# Denoising large-scale biological data using network filters

**DOI:** 10.1101/2020.03.12.989244

**Authors:** Andrew J. Kavran, Aaron Clauset

**Affiliations:** Department of Biochemistry, University of Colorado, Boulder, CO, USA; BioFrontiers Institute, University of Colorado, Boulder, CO, USA; Department of Computer Science, University of Colorado, Boulder, CO, USA; Santa Fe Institute, Santa Fe, NM, USA

## Abstract

Large-scale biological data sets, e.g., transcriptomic, proteomic, or ecological, are often contaminated by noise, which can impede accurate inferences about underlying processes. Such measurement noise can arise from endogenous biological factors like cell cycle and life history variation, and from exogenous technical factors like sample preparation and instrument variation. Here we describe a general method for automatically reducing noise in large-scale biological data sets. This method uses an interaction network to identify groups of correlated or anti-correlated measurements that can be combined or “filtered” to better recover an underlying biological signal. Similar to the process of denoising an image, a single network filter may be applied to an entire system, or the system may be first decomposed into distinct modules and a different filter applied to each. Applied to synthetic data with known network structure and signal, network filters accurately reduce noise across a wide range of noise levels and structures. Applied to a machine learning task of predicting changes in human protein expression in healthy and cancerous tissues, network filtering prior to training increases accuracy up to 58% compared to using unfiltered data. These results indicate the broad potential utility of network-based filters to applications in systems biology.

**Author Summary:** System-wide measurements of many biological signals, whether derived from molecules, cells, or entire organisms, are often noisy. Removing or mitigating this noise prior to analysis can improve our understanding and predictions of biological phenomena. We describe a general way to denoise biological data that can account for both correlation and anti-correlation between different measurements. These “network filters” take as input a set of biological measurements, e.g., metabolite concentration, animal traits, neuron activity, or gene expression, and a network of how those measurements are biologically related, e.g., a metabolic network, food web, brain connectome, or protein-protein interaction network. Measurements are then “filtered” for correlated or anti-correlated noise using a set of other measurements that are identified using the network. We investigate the accuracy of these filters in synthetic and real-world data sets, and find that they can substantially reduce noise of different levels and structure. By denoising large-scale biological data sets, network filters have the potential to improve the analysis of many types of biological data.

## INTRODUCTION

System-wide measurement data, whether molecular, cellular, or ecological, are often contaminated by noise, which can obscure biological signals of interest. Such noise can arise from both endogenous biological factors and exogenous technical factors. In molecular profiling data, factors include reagent and protocol variability, researcher technique, passage number effects, stochastic gene expression, and cell cycle asynchronicity. This variability can mask underlying biological signals when measuring cell state and how it changes under different conditions, e.g., in development [1, 2], cancer progression [3], and adaptive drug resistance [4, 5]. Noise has also been implicated in the appearance of false signals and in the non-replicability of some studies [6, 7]. Identifying and correcting noisy measurements before analysis is likely to improve the detection of subtle biological signals and enable more accurate predictions in systems biology.

If correlations between related molecular signals are stronger than correlations among sources of noise, then distinct but related signals can be combined to denoise biological measurements, at the expense of a smaller effective sample size. There are three common approaches to identifying related signals: gene sets, subspace embedding, and networks. In the first category, methods like GSEA [8, 9] use the enrichment of genes within curated sets to project the data onto biologically relevant features. While gene sets can increase the power to identify differentially regulated processes, they are inherently coarse, and can themselves be noisy, incomplete, or biased, and thus may not generalize to novel processes. Subspace embedding techniques include PCA [10], clustering [11], and neural network autoencoders [12, 13]. These methods can capture novel gene-gene correlations, but they rarely incorporate biological information into the feature extraction, which can limit both interpretability and generalizability.

Molecular profiling data alone does not directly inform which measurements should be more or less related to each other. Networks that represent a molecular system’s functional structure can provide this missing information. For example, protein-protein interaction, metabolic reaction, and gene regulation networks each encode precise and biologically meaningful information about which groups of measured protein expression levels, metabolite concentrations, or transcript levels are functionally related, and hence which measurements should be combined to filter out independent noise.

Among neighboring elements in the network, the underlying signals may be correlated (assortative) or anticorrelated (disassortative) [14]. For example, differential expression tends to correlate between neighboring genes in a regulatory network [15]. In contrast, inhibitory or compensatory interactions [16, 17] will tend to produce a disassortative relationship. Beyond pairs of measurements, networks can also exhibit large-scale mixing patterns among these interactions, such that a network may be more or less assortative in some regions and disassortative in others [18]. Existing network-based methods typically do not exploit this variability, and instead assume globally assortative mixing by applying a single filter to the whole network [19]. Mismatching the filter and the relationship type, e.g., an assortative filter with anti-correlated measurements, can further obscure the underlying biological signals. Here, we describe a general network-based method that can automatically detect large-scale mixing patterns and account for both assortative and disassortative relationships.

These network filters are closely related to kernel-based methods in image processing [20], in which groups of related pixels are transformed together to improve their underlying visual signal. Most such techniques leverage an image’s underlying grid geometry to choose which pixels have related signals for denoising. Networks lack this geometry because a node’s interactions are inherently unordered, whereas the left- and right-hand neighbors of a pixel are clearly defined. This connection between network filters and image processing is rich with potentially useful ideas that could be adapted to process large-scale biological data. For instance, community detection in networks is a clear analog of the common “segmentation” step in image analysis, in which pixels are first partitioned into groups that represent the large-scale structure of an image, e.g., to separate foreground and background, or a car from the street, and then different filters are applied to each segment (module).

We first describe two classes of network filters, which combine measurement values from neighboring nodes to calculate an assortative or disassortative denoised value, and we describe a general algorithm that decomposes the network into structural modules and then automatically applies the most appropriate filter to the nodes and connections within each module. When applied to synthetic data where the true values and network structure are known, these filters substantially reduce errors relative to a baseline. In addition, we show how applying the wrong filter with respect to the underlying biological relationship can lead to increased errors. Finally, to test the practical utility of these methods in a more realistic setting, we investigate the impact of network filtering on a machine learning task in which we predict changes in human protein expression data when a healthy tissue becomes cancerous. Using the network filters to denoise the expression data before model training increases the subsequent prediction accuracy up to 58% compared to training on unfiltered data.

## RESULTS

### Network filters

A network filter is specified by a function *f* [*i*, **x**, *G*], which takes as input the index of the measurement (node) to be denoised, the list of all measurements **x**, and the network structure *G* among those measurements. The output is the denoised value 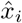. Here, we consider only local network filters, which use the measurement values of *i*’s immediate neighbors in *G*, denoted by the node set *ν_i_*, which are likely to be the most biologically relevant for denoising. Each filter is applied synchronously, so that all denoised values are obtained simultaneously to prevent feedback within the denoising process.

We note that the idea of a network filter can naturally generalize to exploit information, if available, about the sign or strength of interactions in *G*. This information can be encoded by an edge weight *w_ij_*, which can capture inhibitory or excitatory interactions that are strong or weak. Below, we focus on the case in which this information is not available.

When a measurement *x_i_* correlates with the values of its neighbors 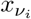 in the network (assortativity), a network filter should adjust *x_i_* to be more similar to the measured values of its neighbors (Fig. 1A). Among the many choices of functions with this qualitative behavior, the mean and median have useful mathematical properties, and connect with past work [19]. This setting is analogous to a smoothing operation in image processing, in which a pixel’s value is replaced by the mean or median of its value and its neighbors’ values. In the context of a network, the mean and median “smoothing” filters have the forms:

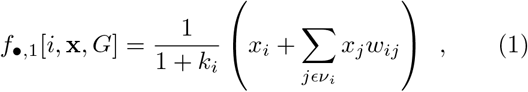

where *w_ij_* = 1 and *k_i_* is the degree of node *i*, reflecting unweighted interactions, and

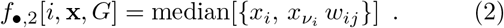

**FIG. 1.**
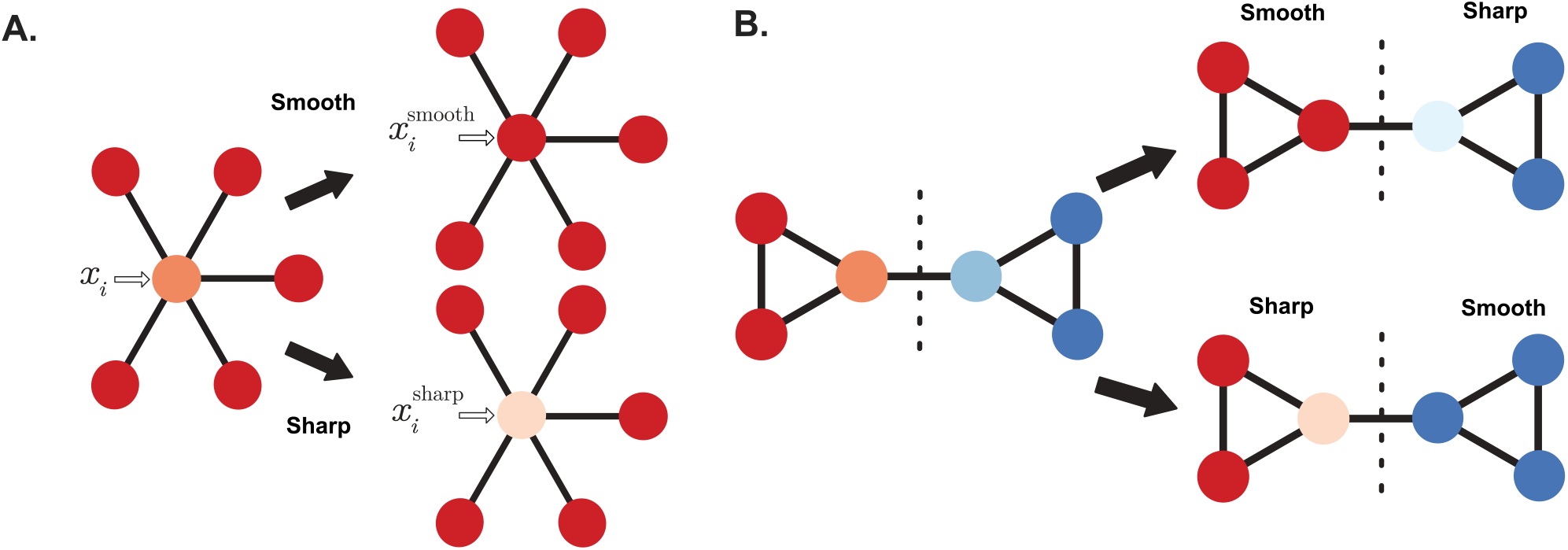
Schematics of Network Filters. Network filters are tools that denoise real-valued biological data using a biologically meaningful network to exploit the correlation (“smoothing”)or anti-correlation (“sharpening”) among neighboring measurements. **A.** A measurement *x_i_* and its neighboring values in network, where the color intensity is proportional to the measured value. In applying the smooth filter, *x_i_* is adjusted to be more similar to its neighbors; in applying the sharp filter, *x_i_* is adjusted to be more distant from its neighbors. **B.** Measurements can also first be partitioned into groups (dashed line) by detecting structural modules within the network, and then different filters applied to different modules, ignoring between-module edges, e.g., if the signals are assortative in some communities and disassortative in others.

When a measurement *x_i_* anti-correlates with the values of its neighboring nodes, a network filter should adjust *x_i_* to be more distant from its neighbors (Fig. 1A). This setting is analogous to enhancing the contrast in an image, e.g., when using the technique of unsharp masking to enhance the high frequency signal in an image to make it sharper. In the context of a network, this “sharpening” filter has the form:

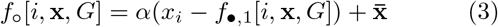

where *α* is a constant scaling factor, and 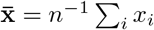 is the global mean. Because *α* is a free parameter, its value should be determined de novo for each data set. For the data sets in this study, we empirically determined the optimal *α* = 0.8 using cross validation.

When a system exhibits large-scale mixing patterns of assortative and disassortative relationships, a network should first be partitioned into structural modules using a community detection algorithm, so that relationships within each module are more homogeneous. Let 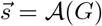 denote the result of applying a community detection algorithm 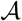 to network *G*, and say that 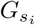 denotes the subgraph of nodes and connections within the module *s_i_* that contains node *i*. Given such a modular decomposition 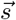, a filter can then be applied to only the subgraph 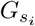 that contains measurement *i*. As a result, relationships that span the boundary between two modules will have no influence on the filtered values (Fig. 1B).

After partitioning, the same filter can be applied to every community, or sharp and smooth filters can be applied to communities with more or less assortative values, respectively. We define such a “patchwork filter” as:

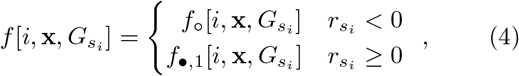

where 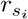 is the standard assortativity coefficient calculated over observed values within community *s_i_* [14]. Any community detection algorithm can be used for 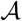. Here, we use the degree-corrected stochastic block model or DC-SBM [21] or the “metadata-aware” version of DC-SBM [22], which are considered state-of-the-art methods [23].

### Tests using synthetic data

We evaluated the performance of these network filters in two controlled experiments with either non-modular or modular synthetic networks, and varying structures and levels of noise.

In the first experiment, we generated simple random graphs with heavy-tailed degree distributions (see Methods) and assigned each node a value drawn from a Normal distribution with mean *μ* = 100 and standard deviation *σ* = 10. These values were drawn in such a way that the assortativity coefficient of the network ranged from *r* ∈ [−0.8, 0.8] (see Methods). As a result, connected values ranged from being highly anticorrelated to highly correlated. To simulate independent measurement noise, we permuted the values among a uniformly random 25% of nodes, and then denoised these “corrupted” values. We find qualitatively similar results for other choices of the fraction permuted. Results report the mean absolute error (MAE) of a denoised value, averaged over 5000 replications.

Without a filter, the average error of a “denoised” value is independent of the underlying correlation (assortativity) among connected values, because this nearby information is left unexploited (Fig. 2A). In contrast, applying a network filter to denoise the corrupted values can substantially improve their accuracy, depending on how strongly coupled a measurement’s true value is with its neighbors’, and what filter is applied to recover that information. For the particular parameters of this experiment, filtering can reduce the error by 37–50% over no filter, and by roughly 20% even in the case of uncorrelated signals (*r* = 0), due to a regression to the mean effect. Error reductions are largest when a network “smoothing” filter is applied to strongly assortative signals, and when a network “sharpening” filter is applied to strongly disassortative signals. That is, denoising works best when the underlying signal structure is matched with the assumptions of the filter.

**FIG. 2.**
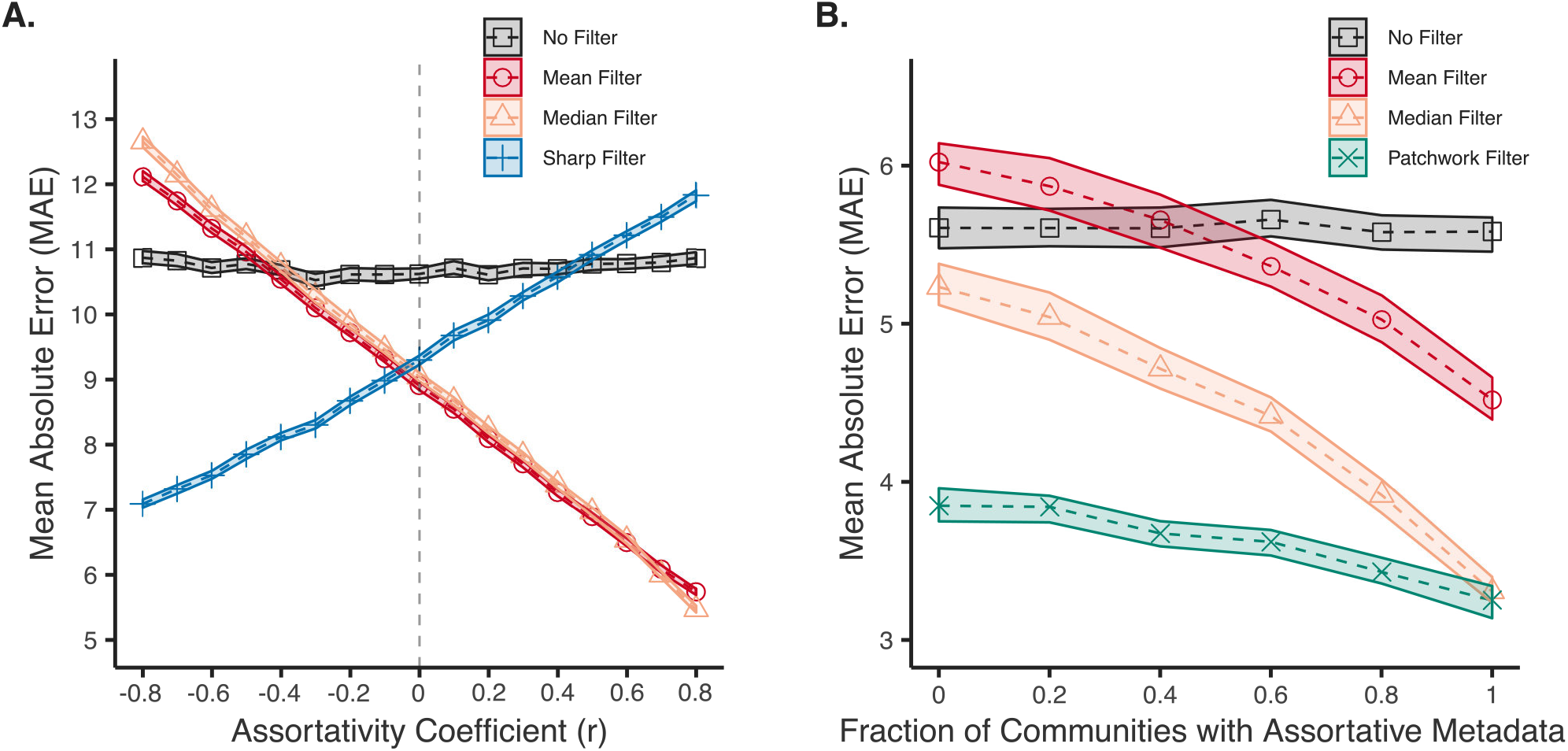
Filter performance on synthetic networks. Network filter tests on synthetic graphs with varying structures and known noise. **A.** The Mean Absolute Error (MAE) of network filters on the permuted nodes as a function of the assortativty coefficient of 5,000 instances of noisy non-modular graphs. The smooth filters (mean and median) perform best on assortative data (*r* > 0), while the sharp filter is optimal for disassortative data (*r* < 0). **B.** The MAE of network filters on the permuted nodes as a function of the fraction of communities with assortative data values for 100 instances of noisy modular graphs. Each network instance has 5 communities and we vary how many communities have assortative vs. disassortative data values with a moderate assortativity coefficient |*r*| ∈ [0.4, 0.7]. The shaded areas indicate 99% bootstrapped confidence intervals.

When the wrong filter is applied, however, error rates can increase relative to not filtering. In such a case, the filter creates more errors in the data than it corrects. On the other hand, this “mismatch” penalty only degrades the overall accuracy at very high levels of correlation (anti-correlation) among the signals, where its magnitude exceeds the natural benefits of filtering at all (Fig. 2A). When the underlying correlations are moderate (|*r*| < 0.4), the average benefits of network filtering will tend to outweigh the average error induced applying the wrong filter.

In the second experiment, we again generated simple random graphs with heavy-tailed degree distributions, but now also with modular structure, which better captures the structure of empirical biological networks (see Methods). These modules denote groups of nodes that connect to other groups in statistically similar ways. For instance, protein interaction networks can be decomposed into groups with similar biological function, and these groups can have distinct types or levels of signal assortativity [18]. In this situation, applying a single filter to all parts of the network could introduce bias in the denoised values, by pooling nearby measurements indiscriminately, compared to filtering modules independently.

Here, we plant *κ* = 5 modules in the same kind of synthetic network as our first experiment, set each module to have a different mean value, and then vary the fraction of modules that have a positive assortativity coefficient |*r*| ∈ [0.4, 0.7] vs. a negative coefficient (see Methods). This kind of signal heterogeneity across modules mitigates the denoising benefits of a simple regression to the mean, and provides a harder test for denoising methods. Given these choices, we generated values within a module, and simulated measurement noise as in the previous experiment (see Methods). In addition to the previous filters, we also apply the “patchwork” filter in this experiment.

As before, the average error of a denoised value with no filter provides a consistent baseline against which we may assess improvements from filtering (Fig. 2B). And similarly, the error for both the smooth and median filters falls steadily as the fraction of modules with assortative signals increases. For the particular parameters of this experiment, the median filter performs roughly 20% better than the mean filter, reflecting the median’s wellknown robustness to outliers, which arise here from the planted signal heterogeneity.

The global sharp filter works poorly for all ratios when applied uniformly across the whole network (Fig. S1). Because each module has a distinct mean value, the global sharp filter generates errors by assuming the global mean is a good representation of the whole network.

In contrast, the patchwork filter is substantially more accurate than any other filter, and exhibits less dynamic range in its error, across different degrees of modular assortativity (Fig. 2B). For the particular parameters of this experiment, the patchwork filter reduces the mean error by 32–40% compared to no filtering, and by 0–37% compared to median or mean filtering. Only when all of the modules are assortative does the median filter match the patchwork filter’s accuracy. This advantage arises because the patchwork filter avoids applying the same filter to different types of underlying signals, if the structure of those signals correlates with the structure of the network (as it does here). That is, applying a single filter to a modular network can introduce errors when denoising, if the local mixing patterns across modules are heterogeneous. Pairing a community detection algorithm with network filters can avoid this problem by identifying large groups of nodes that should be filtered together, in much the same way that different image filters can be applied after first segmenting an image into distinct regions.

### Denoising protein expression levels in cancer

To evaluate the utility of network filters for denoising biological data in realistic settings, we construct a machine learning task in which we predict the precise changes in human protein expression levels when a healthy tissue becomes cancerous (see Methods). This task has potential applications to detecting pre-cancerous lesions [24, 25]. We then quantify the improvement in out-of-sample prediction accuracy when using a network filter to denoise the input expression data before model training, compared to training on unfiltered data.

For this experiment, protein expression data are drawn from the Human Protein Atlas (HPA) [26], which provides large-scale immunohistochemistry (IHC) measurements for over 12,000 human proteins in 20 tissues, each in both healthy and cancerous states. Antibody based methods like IHC are known to be noisy and prone to variation from uncontrolled experimental parameters [27], which makes this data set a realistic example of noisy large-scale biological data. A standard principle component analysis (PCA) of the raw HPA expression data reveals that the first component correlates with variations in tissue type, while the second correlates with differences between tissue state (healthy vs. cancerous) (Fig. 3A). Some tissues, however, change more than others, and the changes are not always in the same direction. Hence, predicting the precise changes represents a useful and non-trivial machine learning task that network filtering may improve.

**FIG. 3.**
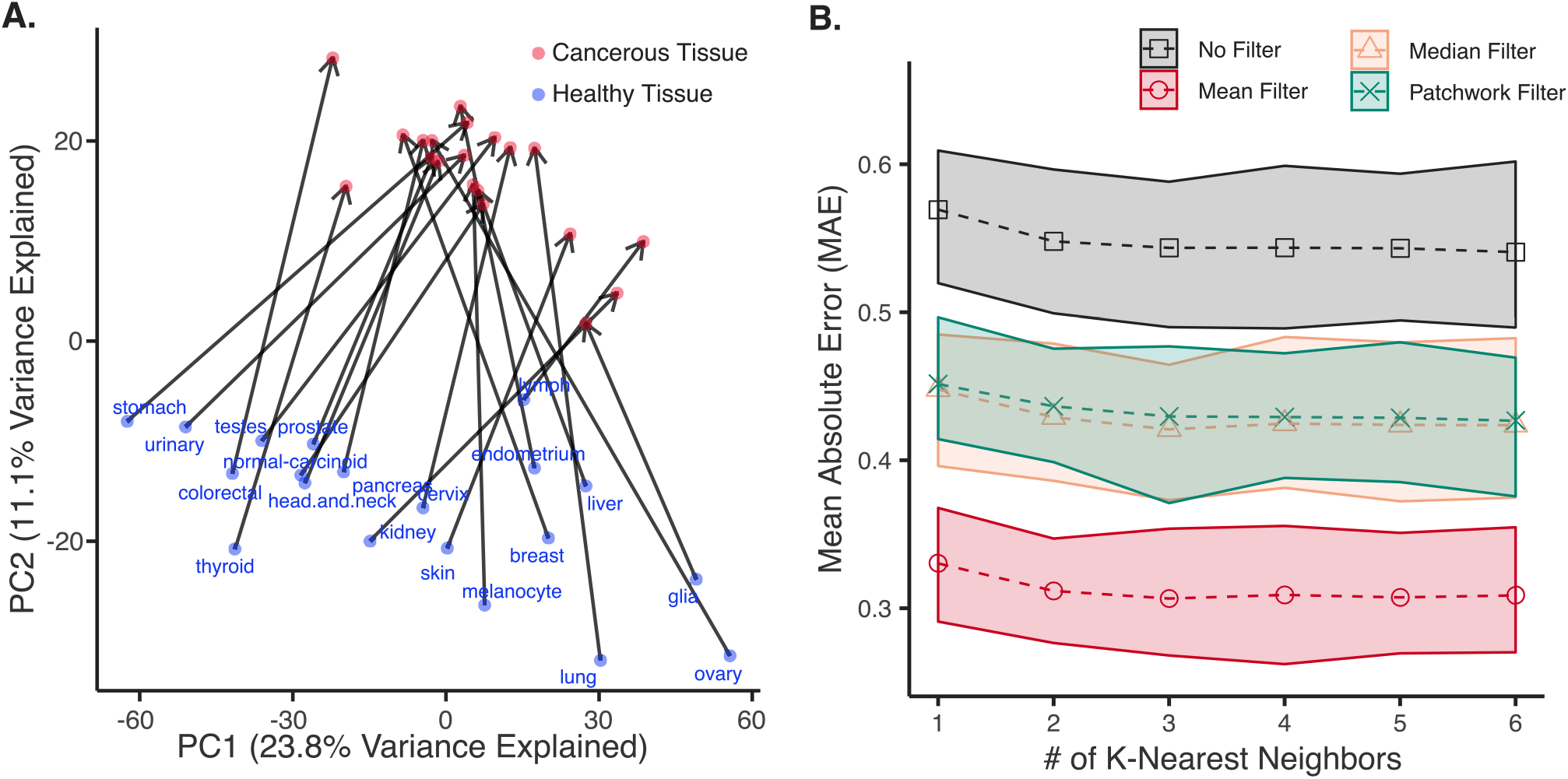
Denoising to Predict Protein Expression Changes in Healthy and Cancerous Tissues. Tests of the network filters on a cancer protein expression prediction task. In this test, we predict the protein expression changes that occur when a healthy tissue becomes cancerous, quantified by the out-of-sample prediction accuracy with and without using network filters to preprocess the data before training. **A.** The first two principal components of immunohistochemistry data of healthy and cancerous tissues in the Human Protein Atlas. Arrows connect a healthy tissue (blue) to the corresponding cancer (red). The first component captures variations across tissues, while the second captures variation in state (healthy vs. cancerous). Predicting the precise changes between healthy and cancerous tissues is a non-trivial task. Therefore, we perform a K-Nearest Neighbors regression on the HPA data, with and without preprocessing with network filters. We evaluate the model by leave-one-out cross validation, and calculating the MAE of the predicted and actual data values for the left out healthy-cancerous pair. **B.** All network filters improve the MAE compared to the no-filter baseline. We compare this across different choices of *K*, as it is a free parameter. The shaded areas represent 99% bootstrapped confidence intervals

For the network filters, we use a comprehensive map of the human protein-protein interaction network (PPIN) [28], which combines data from several interactome databases and is curated for biological interactions with high levels of evidence. While this network represents a broad collection of authoritative interactome data, the completeness of the human PPIN is still uncertain [29], and we do not regard this network as itself noise-free. Taking the intersection of proteins contained in both expression data and interaction network (see Methods) yields data on *n* = 8,199 proteins in a network with *m* = 37,607 edges.

In the machine learning task, we perform a *K*-nearest neighbor regression on an embedded representation of the protein expression data to learn how expression levels change with tissue state (see Methods). We evaluate the trained model via the MAE between the predicted and the actual changes in protein expression under leave-one-out cross validation (in which we train on 19 tissue pairs, and predict on the 20th) with or without denoising the expression data with a network filter prior to model training. Because the number *K* is a free parameter that controls the complexity of the learned model, we evaluate the robustness of our results by systematically varying *K*. For the patchwork filter, we partitioned the graph into 10 modules using the DC-SBM [21] and apply the mean filter within each module. In this data, most measured values are weakly assortative across protein interaction edges, and only a few detected modules exhibit any disassortative signal, and even then their internal *r* is relatively close to zero (Fig. S2). In this situation, the smooth filter typically outperforms the sharp filter (Fig. 2A).

Across model complexities, we find that denoising before model training using any type of smoothing network filter, patchwork or otherwise, provides a substantial reduction in prediction error relative to training on unfiltered data (Fig. 3B, Fig. S3).

Error rates tend to decrease with greater model complexity *K*, suggesting that more complex models are better able to capture variations in the precise expression level changes between tissue states. This decrease in error also occurs without first filtering the expression data. However, the improvement in prediction accuracy from increasing the model complexity without filtering is modest (5.2% at *K* = 6) compared to the improvement from first applying the best network filter (58.2% at *K* = 1, and 57% at *K* = 6).

We note that in this real-world setting, the patchwork filter, which first partitions the protein interaction network into protein groups, performs no better than the simple median filter, and the simple mean filter outperforms both (improving the MAE by 38% at *K* = 6). This behavior suggests that the partition produced by the off-the-shelf community detection algorithm did not correlate sufficiently strongly with the underlying variation in biological signals to correctly localize the most relevant adjacent measurements, in contrast to our controlled experiments (Fig. 2B). Developing community detection algorithms that choose more biologically relevant partitions may be a useful direction of future work.

## DISCUSSION

Large data sets of biological signals, such as system-wide measurements of molecular concentrations, species abundances, or activation levels, are often noisy. However, these measurements are not fully independent because they reflect the dynamics of a single interconnected system. Using a network to represent the underlying biological relationships among a set of measurements, we can leverage the size of these data sets to systematically denoise many measurements at once, improving the data’s utility for understanding the structure and dynamics of complex biological systems or making accurate predictions in systems biology.

These “network filters” are a flexible tool and can exploit a wide variety of network data, including networks of molecular binding interactions, gene regulations, protein interactions, metabolic reactions, cellular interactions, and even ecological species interactions. Network filters can be extended to exploit information about the sign or strength of interactions or to allow the type of interaction to vary across different modules within the network. These filters can also be applied to networks of any size, ranging from local signaling pathways like MAPK or WNT to entire protein interaction networks or connectomes. In fact, any network that correlates with the underlying causal structure of a set of measured variables could potentially be used as a filter. By exploiting these underlying relationships, a network filter pools correlated information, which mitigates independent noise, in much the same way that image processing techniques use information from nearby pixels to denoise an image.

Experiments using synthetic data with realistic biological network structures and a variety of underlying signals indicates that network filters can substantially reduce noise in large biological data sets across a broad range of circumstances (Fig. 2A). The greatest benefit is obtained when the type of filter is matched to the underlying relationship among the signals, e.g., smoothing for assortative signals (correlation) and sharpening for disassortative signals (anti-correlation). However, for modest levels of correlation, even the wrong kind of filter yields some benefit because of a regression to the mean effect, in which combining several related signals filters out more noise than it introduces through bias. When signal types are heterogeneous across the network, so that the strength or direction of the correlation differs in different parts of the network, a “patchwork” filter performs better. In this approach, we first partition the network into smaller, more homogeneous modules (groups of interrelated measurements) and then apply filters independently to the measurements now localized within each module (Fig. 2B).

In a more realistic setting, in which we train a machine learning algorithm to predict changes in human protein expression levels when healthy tissue becomes cancerous, applying a network filter based on a high-quality protein interaction network before model training substantially improves prediction accuracy, compared to training on unfiltered data (Fig. 3B). In this experiment, the protein interaction network itself is not noise-free [29], indicating that filtering using an imperfect network can be better than not filtering at all.

There are a number of potentially valuable directions for future work on network filters, which may improve their error rates or adapt them to more complicated settings or tasks. Techniques from image processing, both simple and advanced, represent a particularly promising direction to explore [30–32]. For instance, here, we only considered filters that combine measurements associated with directly adjacent nodes. As a result, the denoised values associated with low degree nodes in the network derive from a relatively smaller number of measurements, and hence are likely to have larger residual noise than will higher degree nodes. Modifying the network filter for low degree nodes to look beyond nearest neighbors, e.g., to ensure a minimum number of pooled measurements per node, may provide better guarantees on the accuracy of the denoised value. Examples of this type of technique in image processing include the Gaussian filter [33] and methods that weight nodes by personalized PageRank [34] or another random walk method.

Image segmentation, in which an image is first partitioned into visually distinct pieces, e.g., separating the foreground from the background, is a common preprocessing step in image analysis. The patchwork filter considered here is a simple adaptation of this idea, but it relies on an off-the-shelf community detection algorithm to partition the nodes, considers different modules independently, and ignores connections that run between modules. While this approach should retain the most informative relationships among the measurements it also serves to reduce the degrees of many nodes, which may lessen the benefits of filtering, as described above.

Developing filters that utilize the edges between modules could mitigate the induced low-degree effects that come from applying a patchwork filter to account for signal heterogeneity in the system. Such between-module edges should likely be considered separately from within-module edges, e.g., by adjusting their weights *w_ij_* to more accurately capture the character of the particular signal relationship between the modules containing nodes *i* and *j*.

The benefits of a patchwork filter necessarily depends on how closely the network partition correlates with the underlying biological structure of the system. Off-the-shelf community detection algorithms may not always provide such partitions [35], and in some settings, developing application-specific partitioning algorithms, or algorithms that can exploit biologically meaningful node attributes [22], may improve the behavior of a patchwork filter.

Finally, the network filters defined here make few specific assumptions about the underlying noise-generating process itself. In specific applications, much more may be known about the direction, magnitude, and clustering of errors across large-scale measurements. For instance, in molecular profiling data, endogenous biological factors like cell cycle effects likely induce distinct noise patterns compared to exogenous technical factors like sample preparation or instrument variation. Developing more application specific error models that could be combined with network filters may provide more powerful denoising techniques than the general filters described here.

## METHODS

### Synthetic data with known noise and structure

In the first experiment we generate simple non-modular random graphs using the Chung-Lu (CL) model [36–38] with *n* = 100 nodes and a degree distribution that, in expectation, follows a power law distribution Pr(*k*) ∝ *k^−α^* with parameter *α* = 3 for *k* ≥ 1. If the generated degree sequence included a node with degree *k* > 17, a new degree sequence was sampled. This choice ensured that no star-like subgraphs were created. In our analysis, only nodes in the largest connected component were included. This choice mitigates the bias experienced by low degree nodes, which are the most likely nodes to exist outside the largest component.

For each CL synthetic network, we generate node values using the procedure described below. We vary the assortativity coefficient *r* ∈ [−0.8, 0.8] while drawing values from a Normal distribution with mean and variance *μ* = *σ*^2^ = 100. We simulate measurement noise by taking a random permutation of a uniformly random 25% of the node values. We then apply each of the networks filters (mean, median, sharp) to these noisy values, and calculate the mean absolute error (MAE) of the original and denoised values. Results are averaged over 5000 repetition of this process.

In the second experiment, we generate simple modular random graphs using the degree-corrected stochastic block model DC-SBM (DC-SBM) model [21], with *κ* = 5 communities of *nr* = 100 nodes each (*n* = 500 nodes total), and the same degree distribution as the non-modular case. The network’s modular structure is specified using the standard “planted partition” model [21], in which the community mixing matrix *ωrs* is given by a linear combination of a perfectly modular graph and a random graph, and has the form 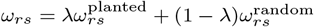, with *λ* = 0.85.

For each DC-SBM network, we generate node values with the following properties: (i) the distribution of values within each module are drawn from a module-specific Normal distribution with mean *μ* = 110, 80, 60, 40, 20 and variance *σ*^2^ = 25, (ii) *κ*′ ∈ [0, 5] communities are assigned to have negative assorativity coefficients, and (iii) the within-community assortativity coefficients are chosen uniformly at random on the interval |*r*| ∈ [0.4, 0.7]. These choices construct a hard test in which a filter’s accuracy is effectively penalized if it uses nodes outside a given community to denoise a particular value. For the patchwork filter, we partition the network using the “metadata-aware” DC-SBM with 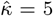 communities [22], using the observed measurements as metadata. Noise is induced and accuracy is assessed as in the non-modular case, except that the nodes are randomly permuted within each module rather than the whole network.

### Generating synthetic correlated measurements

We generate node values with a specified assortativity coefficient *r*_∗_, for a specified adjacency matrix *A*, using Markov chain Monte Carlo (MCMC). The assortativity coefficient *r* is defined as

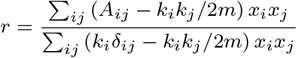

where *k_i_* is the degree of node *i*, *x_i_* is the value associated with node *i*, 2*m* = ∑_*ij*_ *A_ij_* is twice the number of edges in the network, *A_ij_* is the entry in the adjacency matrix for nodes *i* and *j*, and *δ_ij_* is the Kronecker delta function.

Given a network *A*, a desired assortativity coefficient *r∗*, and a node value distribution Pr(*x*), we generate a set of node values follows.

1. Assign each node a value drawn iid from Pr(*x*).
2. Calculate the current assortativity coefficient *r*_0_.
3. Set *t* = 1.
4. While the difference between the desired and current assortativity coefficient Δ = *rt r∗ > β*, a specified tolerance, do:

- Pick a node *i* uniformly at random and assign it a new value 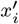 drawn iid rom Pr(*x*).
- Calculate the corresponding assortativity coefficient *r_t_* and difference Δ′ = |*r_t_* − *r*_∗_|.
- If the new value does not improve the assortativity, i.e., Δ′ > Δ restore *x_i_*. Otherwise, increment *t*.
5. Return the node values **x** with the desired assortativity coefficient, *r*_∗_.

In our experiments, we set *β* = 0.009.

### Human protein expression and interaction

Protein expression data were drawn from the Human Protein Atlas (HPA) version 16 [26], which details protein expression in human tissues by large scale immunohistochemistry, for over 12,000 proteins in 20 tissue types, each in a healthy and cancerous state. Human protein interaction (PPIN) data were drawn from the HINT database [28], which combines data from several interactome databases and is curated for biological interactions with high levels of evidence. The HINT network contains *n* = 12,864 proteins and *m* = 62,435 undirected, unweighted edges.

To construct the network filter, we first map the data from the HPA to the PPIN. HPA proteins are indexed by their Ensembl IDs, while HINT proteins are indexed by their Uniprot IDs. A map from Ensembl IDs to Uniprot IDs was constructed using the HGNC BioMart tool. If a node had multiple mapped expression values, we averaged them. We allow protein expression values from HPA to map to multiple nodes if the Ensembl ID maps to multiple nodes in the PPIN. If the gene expression value does not map to any nodes in the PPIN, we discard these as they cannot be de-noised by the network filters. There is one protein in the cancer dataset and 283 proteins in the healthy tissue dataset missing protein expression values in no more than 2 cancers or healthy tissues. For these cases, we impute the missing data from the same protein in another cancer or healthy tissue uniformly at random (impute healthy from healthy, and cancer from cancer).

After keeping the largest connected component of nodes with associated HPA data values, these preprocessing steps produce a network with *n* = 8,199 proteins with IHC expression information across all 20 tissue types and both healthy and cancerous states, and *m* = 37,607 edges. The included healthy-cancerous tissue pairs are: breast, glioma, cervix, colorectal, endometrial, testicular, thyroid, renal, liver, lung, lymphoma, pancreas, prostate, skin, stomach, melanocyte, urinary, head and neck, ovary, carcinoid. For the healthy tissues, the protein expression values of specific cells types that can give rise to the corresponding cancer were averaged together to form one vector (Table S1).

### Predicting expression changes in human cancer

The machine learning task is to predict the changes in protein expression levels when a human tissue changes types from healthy to cancerous. We use *K*-nearest neighbors regression to learn a model that can predict these changes when given the expression levels of a healthy tissue (Fig. 4). We train and evaluate the model using leave-one-out cross validation, in which the model is trained on the observed changes in 19 healthy-cancerous tissue pairs, and is tested on one unobserved pair. We first train and evaluate the model on unfiltered data, and then compare its accuracy to a model where we apply a network filter to the expression data prior to training.

**FIG. 4.**
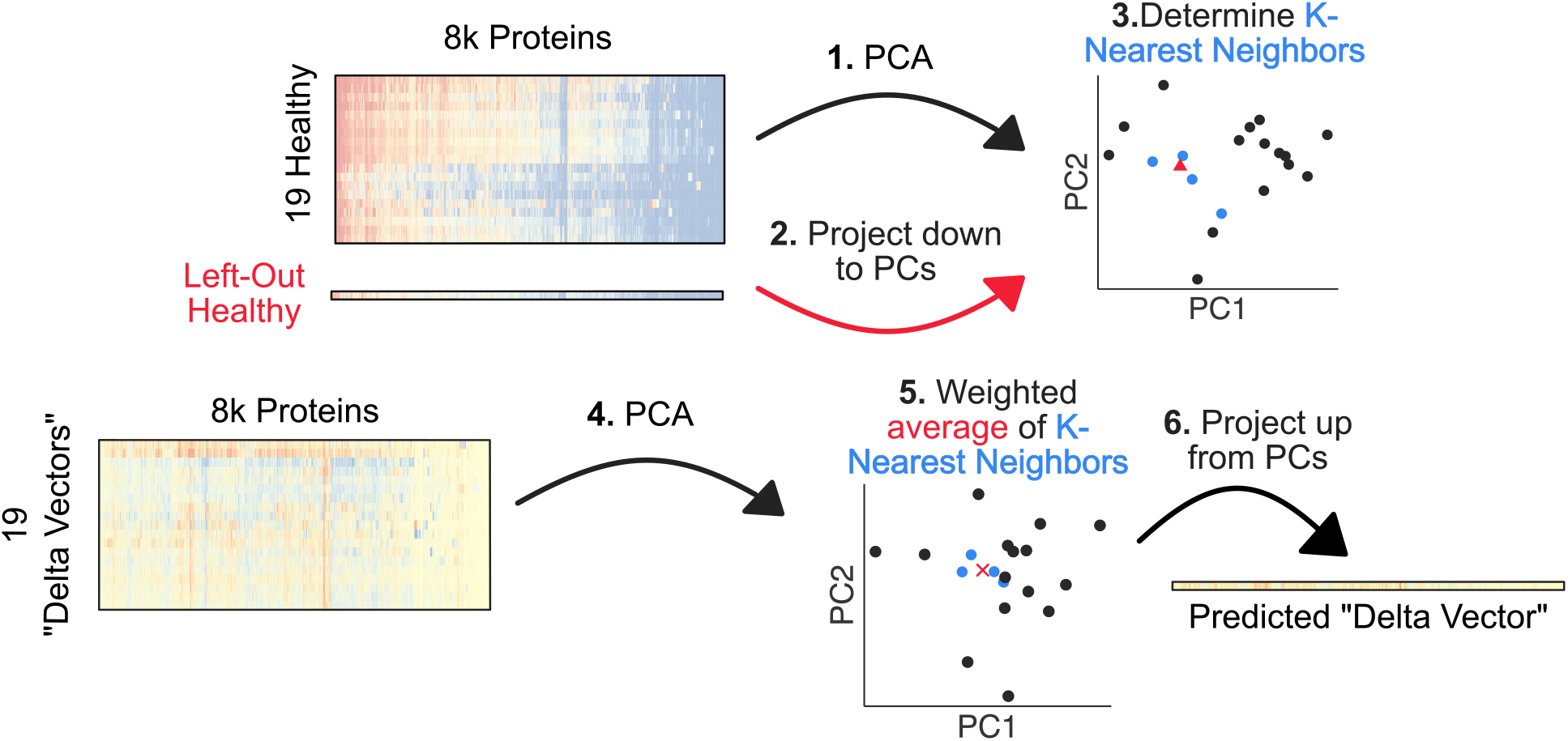
Schematic of K-Nearest Neighbors Regression Framework. We designed a weighted K-nearest neighbors regression framework to predict the protein expression changes a healthy tissue would undergo when becoming cancerous, given a vector of protein expression profile of a healthy tissue. First, we extract features from the training set of 19 healthy tissue protein expression vectors by PCA. Second, we project the left out healthy vector down to the same PCA space, and third, determine K-nearest neighbors to use for the prediction task. Fourth, we extract the features from the 19 delta vectors by PCA, and fifth, predict the delta vector for the left-out healthy sample by taking the weighted average of the K-nearest neighbor’s delta vectors. Finally, sixth, we project the predicted delta vector from PCA space back to a vector of protein expression values to calculate the error.

First, we applied principal components analysis (PCA) on the training set of 19 healthy tissue protein vectors as a feature extraction method. Next, using the embedded PCA space learned from the training set, we project the held-out healthy sample into the same PCA space. We then determine the *K*-nearest neighbors of the held-out healthy tissue by calculating the Euclidean distance of the first four principal components between this point and all other healthy tissues.

Given this identification of which healthy tissues are most similar to the left-out healthy tissue, we predict the protein expression changes for the held-out observation. We calculate the expression changes between cancerous and healthy tissues, which we call a “delta” vector. Then, we perform PCA on the 19 delta vectors to extract features. The weighted average of the delta vectors corresponding to the *K*-nearest neighbors learned from the healthy tissues are averaged together, where the weight is proportional to the inverse of the Euclidean distance to the held-out healthy tissue. Finally, we project the predicted delta vector from four principal components back to the *n* = 8,199 proteins and calculate the mean absolute error (MAE) of this vector and the actual delta vector.

The basic networks filters evaluated in this task have the form given in the main text. For the patchwork filter, we use the DC-SBM to partition the PPIN into *κ* = 10 communities, and apply the mean filter within each community.

## AUTHOR CONTRIBUTIONS

**Conceptualization:** Aaron Clauset, Andrew J. Kavran.

**Data Curation:** Aaron Clauset, Andrew J. Kavran.

**Formal Analysis:** Andrew J. Kavran.

**Funding Acquisition:** Aaron Clauset.

**Investigation:** Aaron Clauset, Andrew J. Kavran.

**Software:** Andrew J. Kavran.

**Visualization:** Andrew J. Kavran.

**Writing - original draft:** Aaron Clauset, Andrew J. Kavran.

**Writing - review and editing:** Aaron Clauset, Andrew J. Kavran.

