## Supplemental Material for "Denoising large-scale biological data using network filters"

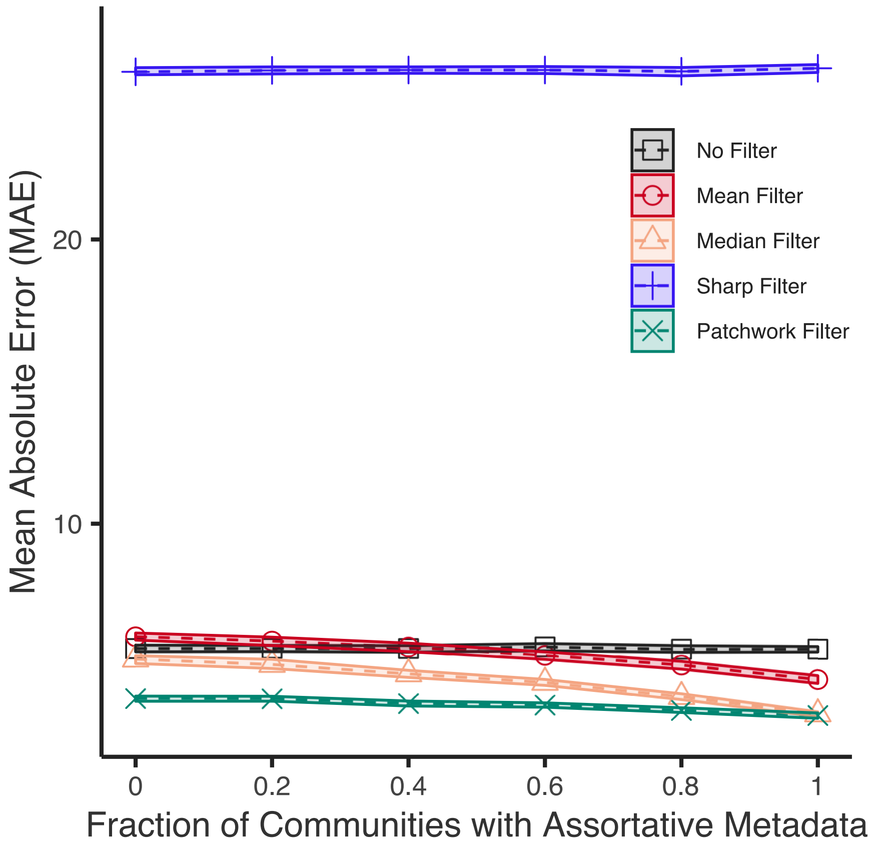


FIG. S1. **Filter performance on modular synthetic networks, including the sharp filter.** The MAE of network filters on the permuted nodes as a function of the fraction of communities with assortative data values for 100 instances of noisy modular graphs. Each network instance has 5 communities and we vary how many communities have assortative vs. disassortative data values with a moderate assortativity coefficient $|r| \in [0.4, 0.7]$. The shaded areas indicate 99% bootstrapped confidence intervals.

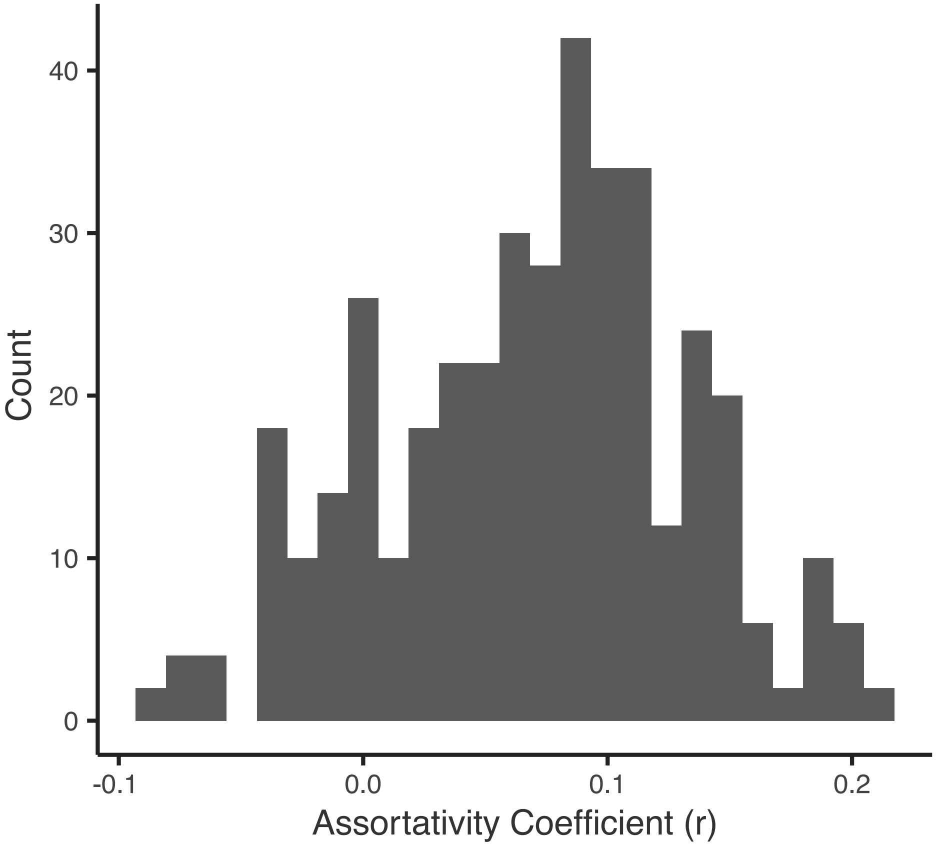


FIG. S2. **Distribution of assortativity coefficients of network modules with Human Protein Atlas Data.** We partition a protein-protein interaction network into 10 modules using the DC-SBM, and map data from the Human Protein Atlas to this network. Then we calculate the assortativity coefficient of each module with the protein expression of 40 different tissue types. Most modules are slightly assortative, while a few are very slightly disassortative.

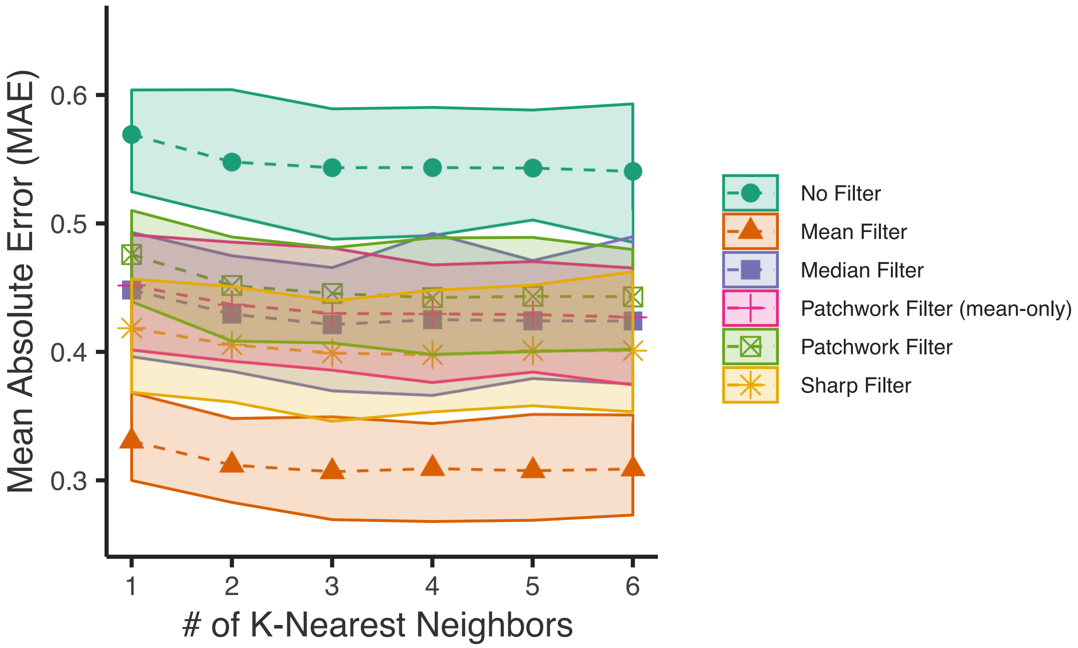


FIG. S3. **KNN regression of Human Protein Atlas data.** We perform a K-Nearest Neighbors regression on the HPA data, with and without preprocessing with network filters. Here we show all simple global network filters in addition to two variations of the patchwork filter: one where we smooth or sharpen within a module depending on the assortativity coefficient, and one where we only apply the mean filter within each module.

TABLE S1. Cell types from the Human Protein Atlas dataset averaged together to form a single healthy tissue vector

| **Healthy Tissue** | **Cell Types Averaged** |
|---|---|
| Breast | Breast Adipocytes, Breast Glandular Cells |
| Glia | Cerebral Cortex Glial Cells, Hippocampus Glial Cells, Caudate Glial Cells |
| Cervix | Cervix Uterine Glandular Cells, Cervix Uterine Squamous Epithelial Cells |
| Colorectal | Colon Endothelial Cells, Colon Glandular Cells, Rectum Glandular Cells |
| Endometrium | Endometrium Cells in Endometrial Stroma, Endometrium Glandular Cells |
| Testes | Epididymis Glandular Cells, Seminal Vesicle Glandular Cells, Testis Cells in Seminiferous Ducts, Testis Leydig Cells |
| Thyroid | Thyroid Glandular Cells |
| Kidney | Kidney Cells in Glomeruli, Kidney Cells in Tubules |
| Liver | Liver Bile Duct Cells, Liver Hepatocytes |
| Lung | Lung Pneumocytes |
| Lymph | Lymph Node Germinal Center Cells, Lymph Node Non-Germinal Center Cells |
| Pancreas | Pancreas Exocrine Glandular Cells, Pancreas Islets of Langerhans |
| Prostate | Prostate Glandular Cells |
| Skin | Skin Fibroblasts, Skin Keratinocytes, Skin Epidermal Cells |
| Melanocyte | Skin Melanocytes |
| Stomach | Stomach Glandular Cells |
| Urinary | Urinary Bladder Urothelial Cells |
| Head and Neck | Nasopharynx Respiratory Epithelial Cells, Oral Mucosa Squamous Epithelial Cells, Salivary Gland Glandular Cells |
| Ovary | Ovarian Stroma Cells |
| Carcinoid (Healthy) | Colon Endothelial Cells, Colon Glandular Cells, Colon Peripheral Nerve Ganglion, Duodenum Glandular Cells, Pancreas Exocrine Glandular Cells, Pancreas Islets of Langerhans, Prostate Glandular Cells |
